# Changes in electrophysiological static and dynamic human brain functional architecture from childhood to late adulthood

**DOI:** 10.1101/2020.05.11.047894

**Authors:** N Coquelet, V Wens, A Mary, M Niesen, D Puttaert, M Ranzini, M Vander Ghinst, M Bourguignon, P Peigneux, S Goldman, M Woolrich, X De Tiège

## Abstract

This magnetoencephalography study aimed at characterizing age-related changes in resting-state functional brain organization from mid-childhood to late adulthood. We investigated neuromagnetic brain activity at rest in 105 participants divided into three age groups: children (6–9 years), young adults (18–34 years) and healthy elders (53–78 years). The effects of age on static resting-state functional integration were assessed using band-limited power envelope correlation, whereas those on transient functional dynamics were disclosed using hidden Markov modeling of power envelope activity. Brain development from childhood to adulthood came with (i) a strengthening of functional integration within and between resting-state networks and (ii) an increased temporal stability of transient (100–300 ms lifetime) and recurrent states of network activation or deactivation mainly encompassing lateral or medial associative neocortical areas. Healthy aging was characterized by decreased static resting-state functional integration and dynamical stability within the visual network. These results based on electrophysiological measurements free of neurovascular biases suggest that functional brain integration mainly evolves during brain development, with limited changes in healthy aging. These novel electrophysiological insights into human brain functional architecture across the lifespan pave the way for future clinical studies investigating how brain disorders affect brain development or healthy aging.

## Introduction

From birth to senescence, humans undergo extensive changes in psychomotor, behavioral and cognitive abilities. These changes are associated with major modifications in structural and functional nervous system architecture, driven by complex interactions between genetic factors and experience.

From childhood to adulthood, the human nervous system encompasses progressive (e.g., neural proliferation, neurite outgrowth, synapse formation) and then regressive (e.g., cell death, axone pruning, synapse elimination) events (for reviews, see, e.g.^1,2^). Progressive events mainly occur during foetal life and set up a broad pattern of neural connectivity, whereas regressive events, which start around birth and end at young adulthood, refine the broad pattern of neural connectivity to a more precise and mature circuitry^1^. Critically, regressive events combine processes that vary in time and space (for reviews, see, e.g.^3–7^). Motor and sensory systems mature before high-order association neocortical areas that integrate those primary functions^3–7^.

From adulthood to senescence, physiological aging is associated with progressive, linear and nonlinear, regional grey matter and more widespread white matter loss, which are due to various processes (e.g., garbage proteins deposits, glial reaction, etc.) eventually leading to neuronal loss, axon elimination, or synaptic density reduction (for a review, see, e.g.^8^).

The advent of structural and functional brain imaging has brought unprecedented insights into the impact of these developmental and aging microstructural processes on long-range functional brain integration. Regressive events characterizing brain development are typically associated with linear and non-linear increase (or strengthening) in functional integration within and between large-scale brain networks (for reviews, see, e.g.^3–7,9^). Physiological aging is mainly associated with a progressive disruption of the established functional integration that mainly involves high-level brain networks, even if inverse processes have also been repeatedly described (for reviews, see, e.g.^9–13^). Still, some studies have suggested that the less functional integration changes are observed through aging, the better is cognitive functioning at old age^14–16^. This suggests that the variability in age-related cognitive and behavioral decline observed in the elders is probably related to individual differences in age-related changes in structural and functional brain architecture.

Age-related changes in human brain functional integration have mainly been investigated using structural and functional magnetic resonance imaging (fMRI) (for reviews, see, e.g.^3–7,10–13^); and much more rarely with positron emission tomography (e.g.^17,18^). Numerous studies relied on task-based fMRI, with the possible confounds of performance bias or reliance on different cognitive strategies between different age groups^9,19,20^. The discovery that human brain activity is organized into resting-state networks (RSNs), i.e., large-scale functional networks active in the absence of any explicit or goal-directed task (for reviews, see, e.g.^21–23^), provided a solution to these critical issues. Although discovered and mostly investigated using fMRI, the electrophysiological equivalent of RSNs were uncovered with magnetoencephalography (MEG)^24–28^ and electroencephalography (EEG)^29–32^ using band-limited power envelope correlation as resting-state functional connectivity (rsFC) index. Compared with fMRI, these electrophysiological techniques have the critical advantage of having an excellent temporal resolution (at the level of the millisecond, for a review see ^33^) and hence can uncover (i) the spectral dynamics of RSNs^24–28^, and (ii) the dynamical aspects (i.e., the spatio-temporal variations) of the functional integration within and between RSNs^25,28,32,34–36^. Moreover, while fMRI relies on an indirect haemodynamic-based measure of brain activity driven by neurovascular coupling, MEG and EEG provide direct information about neuronal activity. This latter aspect reveals also to be critical when it comes to the investigation of age-related brain changes, as age substantially influences the physiological processes at the origin of the neurovascular coupling (for reviews, see, e.g.^20,37,38^). The use of fMRI in the context of age-related investigations therefore introduces a major additional confound, even at rest^38^. MEG and EEG thus appear more appropriate alternatives for such investigations by focusing on direct neuronal activity.

MEG studies that investigated age-related changes in RSNs with *static* rsFC (i.e., estimated over long timescales of several minutes) demonstrated that band-limited power envelope correlations (linearly or nonlinearly) increase from childhood to adulthood within and between RSNs, mainly in the alpha and beta frequency bands^39–41^. They also showed that healthy aging (i.e., selected elder participants without any confounding factors such as vascular disease or cognitive decline) is characterized by the absence of significant changes in functional integration within and between RSNs^16^, supporting the *brain maintenance* theory that proposes that preserved brain architecture contributes to preserved cognitive functioning. Some studies also took advantage of the high temporal resolution of MEG to address age-related effects on the dynamical spatio-temporal variations (i.e., over supra-second or sub-second timescales) in functional integration within and between RSNs^16,41^. One study that investigated these effects in healthy aging using sliding window rsFC (i.e., rsFC estimated over short time windows of prespecified width, typically a few seconds; for a review, see, e.g.^42^) failed to find substantial age-related changes within and between RSNs^16^. Another study investigated fine (sub-second) temporal aspects from mid-childhood to early adulthood using an alternative approach based on hidden Markov modeling (HMM) of MEG power envelopes^41^. Compared to sliding windows rsFC, the HMM identifies transient network configurations (henceforth referred to as “states”) by classifying distinct patterns of envelope (co)variance consistently repeating in time (for a review, see, e.g.^42^), without the need to fix *a priori* the width of a sliding time window. From MEG data, about 6–8 (or more, see, e.g.^43^) transient recurring states lasting 50–200 ms are typically disclosed with spatial network topography quite similar to that of some RSNs^35,43–46^. Using this approach, it has been shown that, as children (>9 years) grow in age, four states mainly encompassing bilateral temporal and parietal cortices exhibit significant nonlinear monotonic age-related decrease (two states) or increase (two states) in state power (i.e., the global, whole-brain change in power envelope that occurs during each state visit)^41^. Among them, one state mainly encompassing bilateral temporo-parietal junctions (TPJs) demonstrated a significant relationship with the spatial signature of static rsFC changes with age. Further, both the time spent in that state on a single visit and the fraction of recording time that the brain spent in that state increased with age. These findings thus suggested that, as the human brain matures, increases in static functional integration of core attentional areas are associated with increased temporal stability within these areas. Of note, the HMM of MEG envelopes has also been used to investigate the impact of pathological aging on transient brain network dynamics^44^, but data on healthy aging or from childhood to late adulthood are, to the best of our knowledge, lacking. Also, previous MEG envelope HMM studies disclosed a transient state with a network topography resembling that of the default-mode network (DMN)^35,44–46^. Critically, this state did not encompass the posterior midline part of the DMN (i.e., the precuneus and the posterior cingulate cortex (PCC)), possibly due to methodological issues related to the type of source reconstruction^47^. These brain areas are of utmost interest when it turns to the investigation of age effects on the human brain architecture considering (i) their critical associative and integrative functions, and (ii) the fact that age may have a specific impact on the functional integration of these brain areas with the rest of the brain^48,49^. It therefore appears critical to investigate age-related effects on transient brain network dynamics using methods better suited to investigate the DMN as a whole^47^.

The present study aimed at characterizing the age-related changes in resting-state functional brain organization, from mid-childhood to late adulthood. To this end, we analyzed resting-state MEG data in 105 participants divided into three age groups encompassing mid-childhood, early and late adulthood. Both static rsFC connectome and HMM state dynamics were investigated using MEG power envelopes (for a detailed description of the value of connectome analyses for lifespan studies, see ^48^). MEG sources were reconstructed via Minimum Norm Estimation (MNE, see below), the method best suited to uncover posterior midline cortices of the DMN^47^. We expected (i) to replicate previous developmental and healthy aging MEG findings about age-related changes in static functional integration within and between RSNs, (ii) that access to the posterior midline cortices of the DMN would bring novel insights into the age-related changes in the static functional integration and dynamic stability of that core human brain network, and (iii) that studying three different age groups from mid-childhood to late adulthood would enhance the understanding of age effects on the electrophysiological brain architecture compared with more classical, separate comparisons of children vs. young adults and young adults vs. elders.

## Results

Five minutes of eyes-open, resting-state MEG activity were recorded in the sitting position using a whole-scalp MEG in 105 healthy participants divided into three groups: 32 children (age range: 6–9 years, mean age ± SD: 7.8 ± 0.9 years), 38 young adults (18–34 years, 23.3 ± 3.8 years) and 35 elders (53–78 years, 66.1 ± 5.9 years). Elder participants were rigorously selected to be considered as *healthy* elders, i.e., participants without any psychotropic drug consumption, sleep impairment, neurologic, psychiatric or cognitive confounding factors.

Static rsFC was investigated first to focus on age-related changes in functional integration, using band-limited power envelope correlation of MNE-reconstructed source activity within 3 frequency bands (theta, θ: 4–8 Hz; alpha, α: 8–12 Hz; beta, β: 12–30 Hz). The functional connectome was built by measuring rsFC among 32 brain regions distributed across 6 well-known RSNs: the DMN, the language, the ventral and dorsal attentional, the primary visual and the sensorimotor networks^16,50^. Envelope correlation was estimated after pairwise signal orthogonalization^51^ and low-pass filtering (1 Hz) of power envelopes. Effects of static power, sex and the MEG system version (Vectorview vs. Triux) used in this study were regressed out of the rsFC data before further analysis. Global connectivity (i.e., mean static rsFC value across all 496 connections) and global power (i.e., mean power across all 32 RSN nodes), as well as mean network connectivity (i.e., mean static rsFC values across all connections within each RSN) and mean network power (i.e., mean power across all nodes of each RSN) were also computed in each frequency band. Group-level (i.e., children, young adults and elders) differences in these summary rsFC and power measures as well as in the detailed rsFC connectome were assessed using non-parametric ANOVA (Kruskal-Wallis tests) with post-hoc Tukey’s range test on ranks to identify age-related effects. Significance was set at *p* < 0.05 with Bonferroni correction for the false positive rate, which is inflated due to multiple testing across three frequency bands, six RSNs for mean network measures, 32 nodes or 496 connections within the functional connectome. In the latter case, the Bonferroni factor for the false positive rate relied on a proper estimation of the independent number of nodes/connections^16,41^ rather than their raw number, avoiding unduly statistical strictness.

Figure 1 depicts global connectivity and global power per age group and frequency band, as well as the significant differences between groups. Figure 2 provides a similar illustration for mean network connectivity and mean network power associated with each RSN.

**Figure 1:**
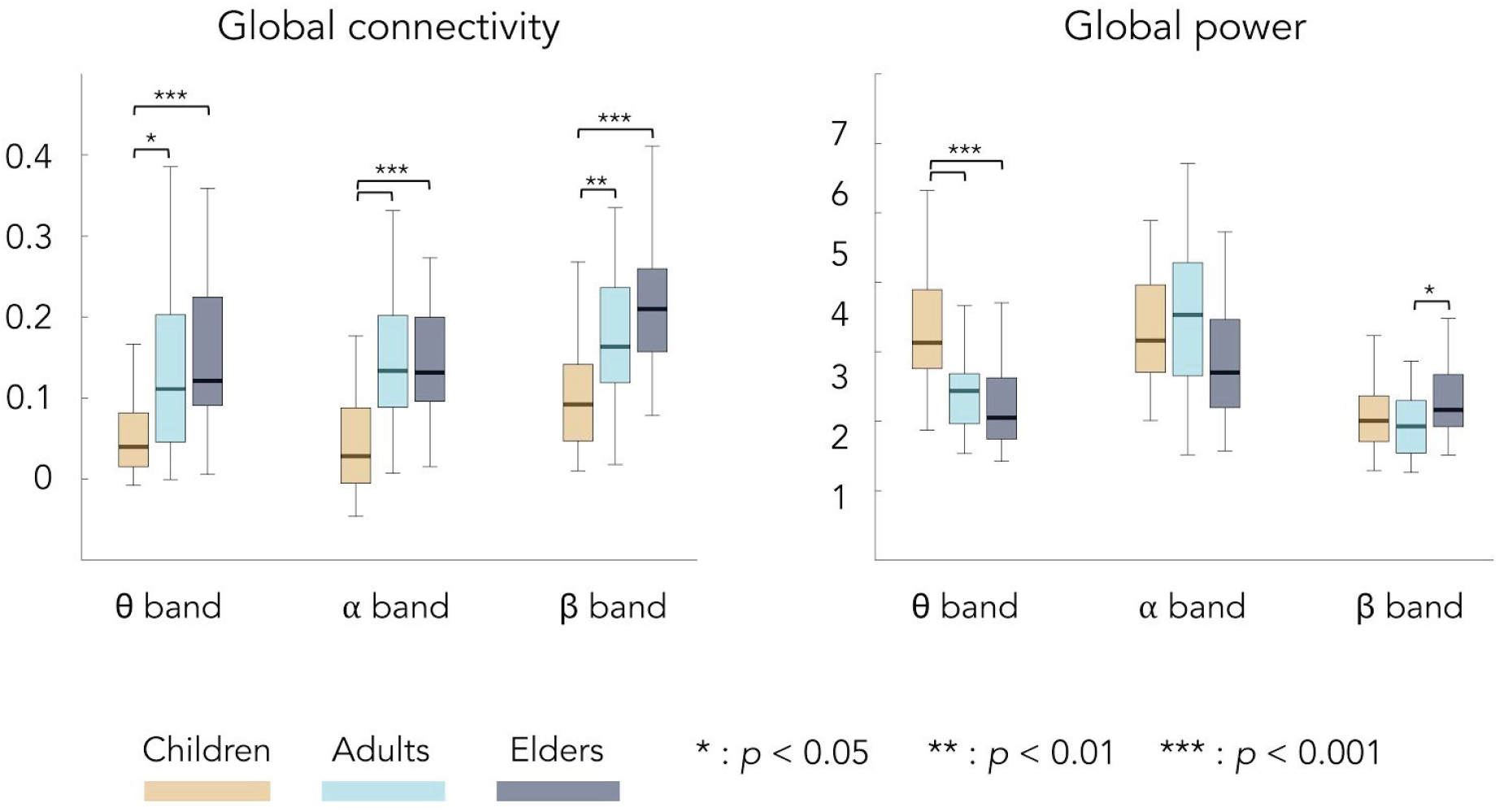
Global connectivity and power for each frequency band and age group (orange, children; light blue, young adults; dark blue, elders). Bottom and top edges of the boxes indicate the 25th and 75th percentiles. Thick middle lines indicate the median. Extreme bars extend to minimum and maximum values (excluding outliers). Statistical differences between groups are indicated with bars along with *p*-values corrected with Bonferroni for 3 comparisons (i.e., the number of frequency bands).

**Figure 2:**
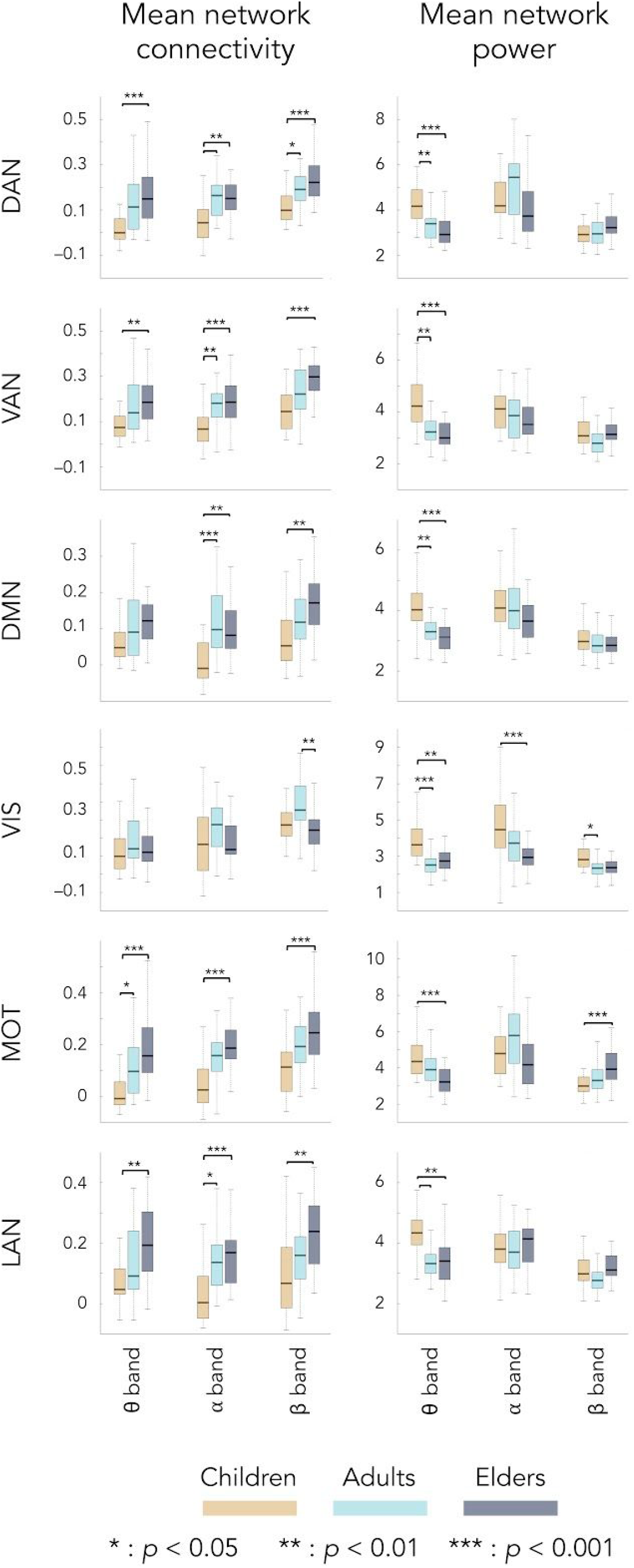
Mean network connectivity and power for each frequency band, age group, and RSN. All is as in Figure 1 except that *p*-values are corrected for 18 comparisons (i.e., three frequency bands times six RSNs).

Global connectivity significantly increased from childhood to early/late adulthood for all frequency bands, with no difference between young adults and elders. Similar age-related changes were observed for mean network connectivity within all RSNs, except for the visual RSN. Mean visual connectivity was unmodulated in the θ and α frequency bands, whereas in the β band it increased from childhood to early adulthood but then decreased in elders back to children’s level. These age-related effects on functional integration were qualitatively different from those on power. The most consistent age-related change on global and mean network power was a significantly higher θ-band power in children compared to adults and elders observed in each and every RSN.

Figure 3 locates the underlying connections within the connectome showing statistically significant age-related rsFC differences, along with the proportion of within- and cross-RSN links involved.

**Figure 3:**
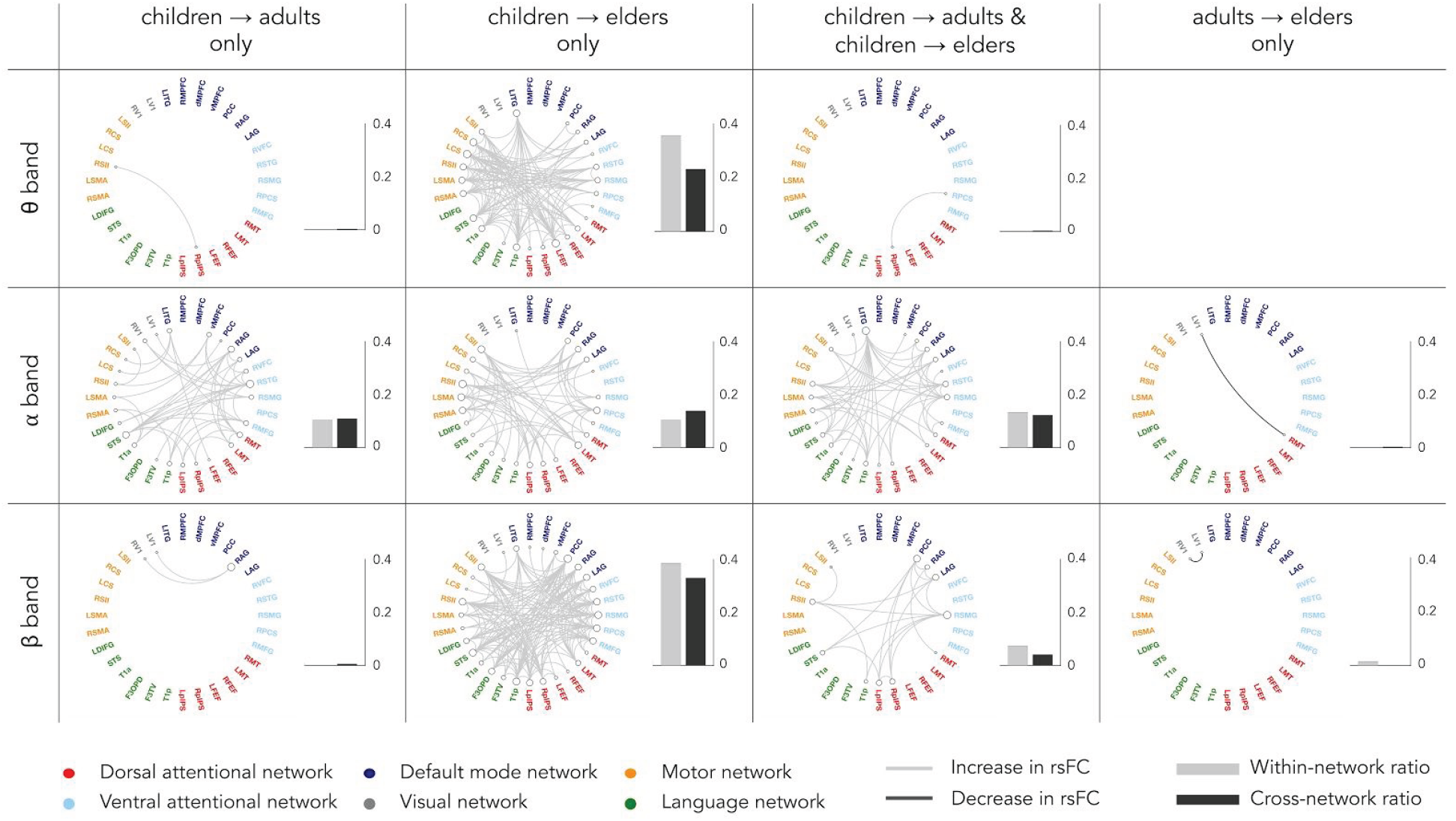
Age-related differences in static rsFC for the three frequency bands and proportion of intra- and cross-RSNs connections showing significant age-related change. Significance was established by post-hoc Tukey’s range test on ranks at *p* < 0.05 corrected for the effective number of band-specific connections (i.e., 513; see Methods). On the circular rsFC plots, light grey lines are related to increase in rsFC, while dark grey lines reveal decrease in rsFC. On the histograms on the right of circular plots, light grey boxes refer to the proportion of within-RSNs connections deemed significant, and dark grey boxes to the proportion of cross-RSNs connections deemed significant.

In line with the summary statistics shown in Figures 1 and 2, static rsFC within and between RSNs increased from childhood to adulthood (young adults or elders) in each frequency band (but mostly in α and β frequency bands). Almost no significant difference was observed between young adults and elders (but two age-related rsFC decreases with the visual RSN). Of note, it is possible that spurious, “ghost” interactions persisting after pairwise orthogonalization^52–54^ contributed to certain connections disclosed in Figure 3. However, the broad rather than localized patterns of rsFC changes encompassing both within and across RSNs cannot be qualitatively altered by ghost interactions, suggesting that these results are robust.

To assess whether these age-related changes in functional integration were accompanied with modifications in dynamic network stability, we used the MEG envelope HMM approach to reduce whole-brain wideband (4–30 Hz) source envelope activity reconstructed with MNE to 8 transient recurrent states^35,44–46^) whose periods of activation/inactivation are determined on the scale of a fraction of seconds (every 25 ms). The HMM along with the Viterbi algorithm returned a binary time series of most probable state activation under the constraint that two states cannot be active simultaneously^55^. These time series allowed us to map the topographical distribution of state power (which measures the degree of regional power increase/activation or decrease/deactivation during state visits). We also extracted temporal parameters characterizing state dynamics such as mean lifetime (MLT, i.e., the mean time spent in each state on a single visit), fractional occupancy (FO, i.e., the fraction of total recording time that the brain spends in each state) and mean interval length (MIL, i.e., the mean time interval between two visits to the same state)^35,41^. Group-level (i.e., children, young adults and elders) differences in these temporal parameters were also assessed using non-parametric Kruskal-Wallis tests with post-hoc Tukey’s range test on ranks. Significance was set at *p* < 0.05 Bonferroni corrected for the number of independent states (i.e., 7).

Figure 4 presents the state power maps (please refer to *Methods* section for a description of the statistical threshold applied) of the 8 HMM transient states. High positive (respectively negative) local state power in a brain area implied that power envelopes in that area tended to increase (respectively decrease) when the brain visited that state.

**Figure 4:**
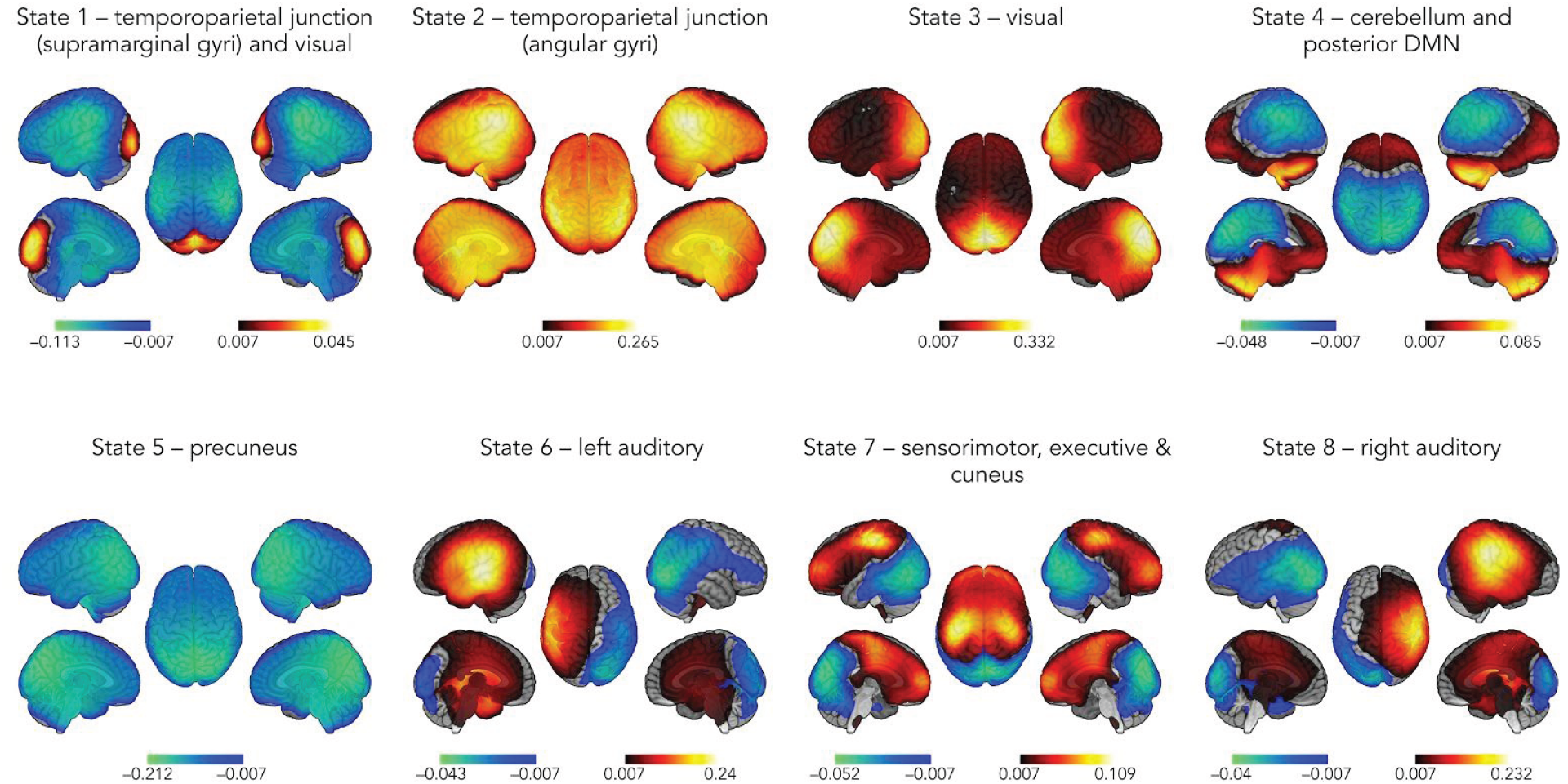
Spatial topographies of the 8 HMM transient states. Red scale refers to the degree of power increase during state visit and blue scale is related to power decrease. These scales are measured in terms of a partial correlation (see Methods).

Topographically, states 1 and 2 both encompassed bilateral TPJs. More specifically, when visiting state 1, power decreased at the supramarginal gyri and increased at the same time in primary visual (V1) cortices. State 2 corresponded more specifically to a power increase at the angular gyri. State 3 was characterized by a power increase at V1 cortices. State 4 showed a combination of power increase at the cerebellum and decrease within the posterior part of the DMN, i.e., precuneus and TPJs bilaterally. State 5 corresponded to a power decrease centered on the precuneus. State 6 combined power increase at the left auditory cortex and decrease at the right extrastriate visual areas, and vice-versa for state 8. State 7 was characterised by a power increase at bilateral sensorimotor areas and prefrontal cortices together with a power decrease at bilateral cunei.

Figure 5 displays the temporal parameters assessing the transience and stability of these states.

**Figure 5:**
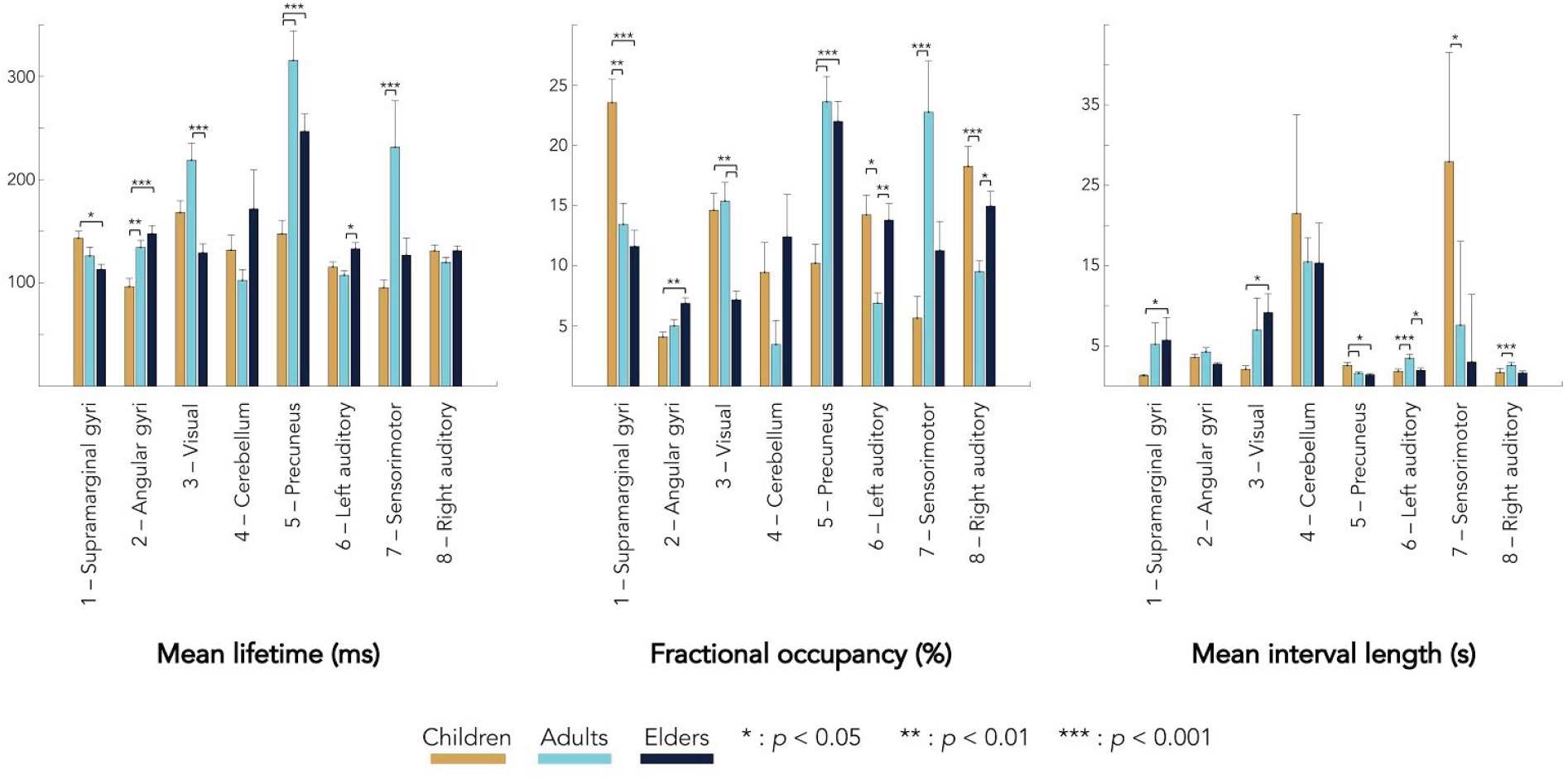
Mean and standard error of mean lifetime (**left**), fractional occupancy (**center**) and mean interval length (**right**) associated to each transient state and age group (orange, children; light blue, young adults; dark blue, elders). Statistical differences between groups are represented by bars along with *p*-values bound on the post-hoc Tukey’s range test on ranks. Here *p*-values are Bonferroni corrected with factor 7 (i.e., number of temporally independent states).

Overall, MLT across all states varied between 100 ms and 300 ms, which is in line with previous MEG envelope HMM studies^35,44–46^. Also consistent with prior studies is that MIL for all states but state 4 ranged between 2 and 6 s in young adults. Surprisingly, the cerebellar/posterior DMN state 4 had substantially longer MIL (15 s). Although this state was visited in all subjects but two, it exhibited considerable gaps between two successive visits, which explains this longer MIL. The posterior DMN associated with this state was partly uncovered in other studies^35,41^ (bar the precuneus, presumably because of methodological issues linked to the source reconstruction method used^47^) but the cerebellum pattern observed here was not present in previous works because cerebellar sources were not modelled. For most states, MLT and MIL appeared inversely related, i.e., longer MLT was associated with shorter MIL and vice-versa. As a consequence, FO (which increases with MLT at fixed MIL, and decreases with MIL at fixed MLT) tended to behave qualitatively as MLT but with sharper variations across groups and states. Accordingly, age significantly affected 7 of the 8 identified states, with state 4 showing no significant age-related effect. For most states (i.e., except for state 2), age tended to affect MLT and FO in similar ways (i.e., increase or decrease), while MIL was affected in opposite ways (i.e., increase when MLT and FO decreases, and vice-versa), as explained above. This means that mainly two types of age-related modulations were observed: either state stabilisation (i.e., increased MLT/FO) with shorter recurrence time (i.e., decreased MIL), or state destabilisation (i.e., decreased MLT/FO) with longer recurrence time (i.e., increased MIL).

States 1 and 2 showed opposite trends of evolution with age for MLT, FO or MIL. A progressive decrease in MLT and FO associated with an increase in MIL, was observed from childhood to late adulthood for state 1, while a progressive increase (but with no effect on MIL) was observed for state 2. These findings indicate that, as subjects age, they visit less often and for a shorter duration a network state in which V1 cortices are active and supramarginal gyri are deactivated, while they visit more often and for a longer duration a network state with active angular gyri. Notwithstanding the involvement of V1, these two opposite effects thus lead to the same conclusion of a progressive stabilisation of power activation in the TPJs from mid-childhood to adulthood. State 3 was dominated by a decrease in MLT and FO from early to late adulthood, with an increase in MIL from childhood to late adulthood. As adults age, V1 activation thus becomes less stable, which is in accordance with the visual part of state 1. State 5 exhibited significantly higher MLT and FO with smaller MIL in adults and elders than in children. Thus, children spend less time in a transient network state of deactivated precunei. In other words, brain development stabilizes precuneus deactivation. State 7 showed significant increases in MLT and FO and a significant MIL decrease from childhood to early adulthood, suggesting the stabilization with brain development of combined sensorimotor and prefrontal activation and cuneal deactivation. Of note, this heightened stability of cuneal deactivation concurs with the destabilization of V1 activation shown by states 1 and 3. The same trends were observed between children and elders without reaching significance. Finally, states 6 and 8 showed similar trends of age-related changes that were predominated by an inverted U-shape from childhood to late adulthood in FO and MIL. These findings showed that young adults spend less time than children and elders in transient network states characterized by an activation of unilateral auditory cortices together with a deactivation of contralateral extrastriate visual cortices.

## Discussion

This study mainly shows that brain development is associated with increased static within- and cross-RSNs functional integration and a dynamic stabilization of power activations in lateral temporo-parietal regions and of power deactivations in midline posterior cortices (i.e., precunei). By contrast, healthy aging was mainly associated with changes in static functional integration and in dynamic stability limited to the visual network.

Results of static rsFC analyses are in line with previous structural and fMRI neuroimaging studies, which demonstrated that microstructural and functional changes accompanying brain development are typically associated with an increase or strengthening in functional integration within and between large-scale brain networks (for reviews, see, e.g.^3–7,9^). Critically, they also parallel those of previous MEG studies relying on envelope correlation as rsFC index^40,41^. Indeed, those studies demonstrated that brain development is characterized by a significant increase in global, mean network, within- and between-RSNs rsFC mainly in the α and β frequency bands from mid-childhood to adulthood. These MEG findings therefore suggest that regressive events together with white matter maturation (i.e., increase in myelination, changes in axonal diameter, etc.) at the core of brain development promote large-scale functional integration through an increase in resting-state neural synchrony^41,56^. Furthermore, the almost total absence of static rsFC changes from early to late adulthood confirms previous results from our group but in a larger (35 instead of 25) sample of healthy elders^16^. The present data provide additional support to the *brain maintenance* theory for successful aging, i.e., the proposal that preserved resting-state functional brain architecture contributes to preserved cognitive functioning. Interestingly, age-related differences in static rsFC from childhood to adulthood were higher when comparing children with healthy elders than with young adults. This was probably related to the relatively low age of young adults (mean age ± SD: 23.3 ±3.8 years), as previous MEG studies suggested that static rsFC relying on amplitude correlation continues to rise after the age of 25 years^39–41^. Additionally, global and mean network θ-band power was also significantly higher in children compared to young adults and elders, which is also a typical finding along brain development^39–41^. As a whole, these static rsFC and power data replicate previous MEG findings and suggest that our participants are representative of the corresponding population for the investigations of age-related changes in transient resting-state brain dynamics.

Using the HMM approach, 8 transient recurrent states were disclosed in our population with rather similar spatial and temporal patterns to those previously described^35,41,44–46^. Importantly, we also found one state (state 5) involving bilateral precunei that was not disclosed in previous MEG HMM studies. This is presumably related to the use of MNE rather than beamforming for source reconstruction in our study, as it was shown that the former is better suited to image midline posterior cortices in functional integration studies^47^. Significant age-related modulation (increase or decrease) in the time spent at rest in 7 out of 8 of those transient recurrent brain states were observed from mid-childhood to adulthood. Some age-related changes appeared very similar to those reported in Brookes *et al*.^41^. In particular, the opposite effects of brain development observed between states 1 (i.e., decrease in the time spent in deactivated bilateral supramarginal gyri) and 2 (i.e., increase in the time spent in activated bilateral angular gyri) concur with their previous finding of stabilization in the power activation of bilateral lateral temporo-parietal cortices. They further suggest that, as children age, the increase in static functional connectivity in these core associative inferior parietal areas is associated with increased temporal stability^41^. These findings were initially attributed to the possible maturation of attentional brain areas^41^. But, considering the many (low- and high-level) cognitive functions supported by those inferior parietal areas and their contribution to many neural networks (for reviews, see, e.g.^57,58^, the functional relevance of these age-related changes) is probably more complex than previously evaluated. Young adults also spent more time than children in transient states of activated sensorimotor and prefrontal (state 7), and less time in transient states of activated left/right auditory and deactivated right/left extrastriate visual networks. Critically, this study also disclosed that brain development was associated with an increase in the time spent in a state (state 5) of deactivated precunei. Taken together, these HMM data suggest that the progress from childhood to adulthood is associated with a maturation of the resting-state transient brain dynamics characterized by an increase in the temporal stability of (i) transient activated networks encompassing associative frontal, inferior parietal and sensorimotor neocortical regions, and (ii) transient deactivation of the precunei. These age-related changes might relate to the previously described developmental increase in the segregation between the precuneus and fronto-parietal networks at rest^59^. The dissociation of the precuneus from the rest of the DMN in a specific deactivated transient state is also probably in line with the recognized DMN functional-anatomic fractionation^60,61^.

From young to late adulthood, this study disclosed that elders spend less time than young adults in a transient state of activated visual network (state 3), and more time in transient state of activated left/right auditory cortex and deactivated right/left extrastriate visual cortices (states 6 and 8). Interestingly, these age-related changes were associated with a significant decrease in static rsFC between left and right primary visual cortices. These findings therefore suggest that healthy ageing is mainly associated with a decrease in static resting-state visual network functional integration and a destabilization of its activations. They may be related with the previously described reduction in the efficiency of occipital visual areas during visual processing^62^ and in the neural specialization of extrastriate visual areas^63^ between young adults and healthy elders. As such, these results obtained in healthy elders contrast with the available fMRI literature that highlighted substantial changes in static functional connectivity with aging (see, e.g.^64–67^). This discrepancy might be due to two possibly interrelated factors^16^. First, we used an imaging method free of neurovascular bias, which is not the case of fMRI. Second, we concentrated on elders with healthy ageing, which may induce the possible drawback that the included elder subjects may actually be considered as not being representative of “typical” elders^16^. This study therefore provides additional evidence highlighting the critical need to compare fMRI and MEG rsFC changes with age in the same population of subjects and in elder subjects with different behavioral and cognitive profiles to better understand the origin of this discrepancy.

A key issue associated with these findings is to determine whether these age-related changes in transient resting-state brain dynamics are linked to the effects of age on spontaneous cognitive processes. Indeed, whether spontaneous dynamic brain activity and functional integration reflects intrinsic (i.e., task-independent) neural processes (e.g., maintenance of homeostasis or the integrity of anatomical connections) or extrinsic (i.e., task-dependent) neural processes, or both, remains an open question (for a review, see, e.g.^68^). As certain subtypes of spontaneous cognitive processes are detectable in time-varying functional connectivity measurements^68^, it could be hypothesized that part of our results might pertain to the age-related changes in the occurrence of mind wandering episodes and in the content/type of spontaneous cognitive processes observed from childhood to young adulthood and from young to late adulthood^63,69–77^. Further studies should investigate this critical issue.

The present study suffers from several inherent limitations. First, we investigated age-related effects on static and dynamic functional brain integration using three different age-groups (children, 6–9 years; young adults, 18–34 years; elders, 53–78 years) rather than a large group of subjects equally distributed from 6 to 80 years. The ensuing between age-group comparisons intrinsically limit the characterization of the observed age-related effects (e.g., linear vs. non-linear effects, critical age(s) for changes, etc.). Second, we did not include children aged under 6 years as they are difficult to measure using conventional cryogenic MEG systems. At that age, many brain systems (e.g., sensory and motor systems) are already largely mature, which means that this study missed part of the maturation of the low-level brain systems. Third, we used a cryogenic MEG system that has been shown to underestimate the level of frontal functional integration due to inhomogeneities in the MEG sensor-brain distance^32,68^ (i.e., in the sitting position, posterior and upper MEG sensors are closer to the brain than anterior sensors). We may therefore have missed or underestimated some age-related changes that occur in anterior brain areas due to a lower signal to noise ratio. Furthermore, the use of a MEG system with fixed helmet size renders the acquisition of high quality data and whole head coverage in young participants more challenging due to their reduced head size. Based on those latter considerations, further studies should rely on on-scalp neuromagnetometers such as optically pumped magnetometers, which have been demonstrated to be usable for lifespan neuromagnetic investigations^78^. Finally, we used a restricted, low-density connectome limited to major RSNs (to limit as much as possible the multiple comparison issue) to investigate the age-related changes in static functional integration. This approach was motivated by the desire to relate the observed changes to mean network, within and cross-RSNs functional connectivity, which is more difficult to operate with a whole brain source-level approach or with a precise parcellation of the human brain such as the automated anatomical labelling atlas^79^. Of note, this goal also argues for the relatively limited impact of orthogonalization asymmetry^52–54^ on our results. Still, this approach probably underestimated the extent of age-related changes in static functional integration.

In summary, this study indicates that brain development combines the strengthening of within and cross-RSNs functional integration with substantial changes in transient resting-state brain dynamics leading to an increase in the temporal stability of (i) transient activated networks encompassing associative frontal, inferior parietal and sensorimotor neocortical regions, and (ii) transient deactivations of the precunei. It also highlights that healthy aging is mainly associated with a decrease in static resting-state visual network functional integration and its temporal stability. As a whole, these results provide novel electrophysiological insights into the effects of age on human brain functional integration from mid-childhood to late adulthood. They also pave the way for future clinical studies investigating how brain disorders can affect brain development or healthy aging.

## Methods

### Participants

Thirty-two children (17 females, mean age±standard deviation (SD): 7.8±0.9 years, range: 6–9 years), 38 young adults (24 females, mean age±SD: 23.3±3.8 years, range: 18–34 years) and 35 elders (24 females, mean age±SD: 66.1±5.9 years, range: 53–78 years) were included in this study. All participants were right-handed according to the Edinburgh handedness inventory (except for one elder who was left-handed), had no prior history of neurological or psychiatric disorder and did not take any psychotropic drugs. No elder reported any subjective sleep or cognitive (e.g., memory impairment) problem, and all had an active personal and social life, did not take any psychotropic drug and were thoroughly screened for sleep habits, depression, anxiety and objective signs of pathological cognitive decline. Based on this comprehensive screening, all elders were considered as *healthy* elders. Twenty elders were included in a previous study from our group and their screening results can be found in Coquelet *et al*.^16^. The other fifteen elders were screened for depression with the Geriatric Depression Scale^80^ (mean scores±SD: 2.4±3.6, range: 0–12), dementia using the Clinical Dementia Rating^81^ (null for all participants) and global cognition with the Mini-Mental State Examination^81–83^ (mean scores±SD: 28.9±0.8, range: 28–30). They also underwent a comprehensive neuropsychological evaluation in which (i) visuoconstructive abilities were assessed using the Rey-Osterrieth complex figure^84^, (ii) cognitive flexibility with the verbal fluency test^85^, (iii) visual episodic memory with the Doors and People Test^86^ (only the Doors part was administered), (iv) working memory using Forward and Backward Digit span^87^, (v) verbal episodic memory from the Free and Cued Selective Reminding Test^88^, (vi) langage oral assessment with Bachy Denomination Test^89^, and (vii) executive functions with the Trail Making Test^90^ (parts A and B) and the color-word Stroop Test^91^. All tests were within the normal range.

Each participant contributed to the study after written informed consent. The CUB Hôpital Erasme Ethics Committee approved this study prior to participants’ inclusion. All experiments were performed in accordance with relevant guidelines and regulations. For each minor participant, written informed consent was obtained from the child (information sheet and informed consent adapted to the child’s age) and one legal representative.

### Data acquisition

Neuromagnetic activity was recorded during 5 minutes at rest (eyes opened, fixation cross, band-pass: 0.1–330 Hz, sampling frequency: 1 kHz) with a 306 whole-scalp MEG system installed in a light-weight magnetically shielded room (Maxshield™, Elekta Oy, Helsinki, Finland; now MEGIN; see ^92^ for detailed characteristics). Ten children, 18 adults and 20 elders were scanned with the Vectorview™ version of the system (Elekta Oy, Helsinki, Finland), while 22 children, 20 adults and 15 elders were scanned with the Triux™ version (MEGIN, Helsinki, Finland) due to a system upgrade. The two neuromagnetometers have identical sensor layout (i.e., 102 magnetometers and 102 pairs of orthogonal planar gradiometers) but differ in sensor dynamic range. Of note, previous works from our group mixing recordings from these two systems did not reveal significant changes in data quality^93,94^, including for static rsFC^32,95^.

In all subjects, four coils continuously tracked their head position inside the MEG helmet. Coils’ location and approximately 200 scalp points were determined with respect to anatomical fiducials using an electromagnetic tracker (Fastrack, Polhemus, Colchester, Vernont, USA).

Participant’s high-resolution 3D T1-weighted cerebral magnetic resonance images (MRIs) were acquired on a 1.5 T MRI scanner (Intera, Philips, The Netherlands).

### Data preprocessing

The signal space separation method^96^ was applied off-line to the continuous MEG data to reduce external magnetic interference and correct for head movements. Then, ocular, cardiac and system artifacts were eliminated using an independent component analysis^97^ (FastICA algorithm with dimension reduction to 30 components, hyperbolic tangent nonlinearity function) of the filtered data (off-line band-pass filter: 0.1–45 Hz). The components corresponding to artifacts were identified by visual inspection and regressed out of the full-rank data.

For source reconstruction, MEG forward models were computed individually on the basis of the participants’ MRI, segmented beforehand using the FreeSurfer software (Martinos Center for Biomedical Imaging, Massachusetts, USA). The MEG and MRI coordinate systems were co-registered using the three anatomical fiducials for initial estimation and the head-surface points to manually refine the surface co-registration (Mrilab, Elekta Oy, Helsinki, Finland). Afterwards, a volumetric and regular 5-mm source grid was built using the Montreal Neurological Institute (MNI) template and non-linearly deformed onto each participant’s MRI with the Statistical Parametric Mapping Software (SPM12, Wellcome Centre for Neuroimaging, London, UK). Three orthogonal dipoles were then placed at each grid point. The forward model model associated with this source space was computed using a one-layer Boundary Element Method as implemented in the MNE-C suite (Martinos Center for Biomedical Imaging, Massachusetts, USA).

### Static resting-state functional connectivity

Cleaned MEG data were filtered in the theta (θ band: 4–8 Hz), alpha (α band: 8–12 Hz) and beta (β band: 12–30 Hz) frequency bands. Band-specific MNE^98^ was applied to reconstruct sources of band-limited activity using planar gradiometers only. The noise covariance matrix was estimated from 5 minutes of empty-room data filtered in the relevant frequency range, and the regularization parameter was estimated from the consistency condition as derived in Wens *et al*.^54^. For power estimation, the depth bias was corrected by a noise normalization scheme, i.e., dynamic statistical parametric mapping^98^. Three-dimensional dipole time series were projected on their direction of maximum variance, and the analytic source signals were then extracted using the Hilbert transform. The functional connectome was restricted to rsFC within a subset of brain regions included in major RSNs (as defined by an fMRI meta-analysis and used in, e.g.^25,50^). This allowed for the investigation of within- and cross-RSNs age-related changes in static functional integration. To that aim, 32 regions of interest were taken from six well-known RSNs (MNI coordinates taken from de Pasquale *et al*.^50^. Specifically, 6 nodes were located in the dorsal attention network, 5 in the ventral attention network, 7 in the DMN, 2 in the visual network, 6 in the motor network and 6 in the language network. Of note, compared to de Pasquale *et al*.^50^, the visual network was restricted to left and right area V1. The resulting rsFC connectome matrices were computed from pairwise correlations of 1-Hz low-pass filtered envelope between each node signals with the others, corrected beforehand for spatial leakage using pairwise orthogonalization^51^. Note that as slight asymmetries might be induced by leakage correction, the resulting rsFC matrices were symmetrized afterwards by averaging it with its transpose. Of note, this approach is suboptimal compared to rsFC computed with inherently symmetric multivariate orthogonalization^52^ and leaves the possibility of remnant “ghost” interactions^53,54^. Source power, estimate as source signals’ variance, was also computed at each node. Finally, global connectivity (i.e., mean connectivity across all connections) and mean network connectivity (i.e., mean connectivity across all intra-RSN connections) were also extracted. A similar analysis was conducted for global power (i.e., mean power across all nodes) and mean network power (i.e., mean power across all intra-RSN nodes).

### Hidden Markov Model dynamic analysis

Sources were reconstructed as in the static approach described hereinabove except that the cleaned data were now wide-band filtered (4–30 Hz). For the HMM analysis of MEG signals, we thoroughly followed the pipeline described elsewhere^35,41^ and implemented in GLEAN (https://github.com/OHBA-analysis/GLEAN), except for the use of MNE as inverse model rather than beamforming. More specifically, source envelopes were computed and downsampled at 10 Hz using a moving-window average with 75% overlap (100 ms wide windows, sliding every 25 ms), leading to an effective downsampling at 40 Hz. Individual datasets of source envelope signals were demeaned and normalized by the global variance across all sources, and then temporally concatenated across subjects. Group-concatenated envelopes were pre-whitened and reduced to their 40 principal components. The HMM algorithm^55,99^ was then run 10 times (to account for different initial parameters and retain the model with lowest free energy) along with the Viterbi algorithm to determine temporally exclusive states of power envelope covariance patterns. We set the number of transient states (*K*) to 8 for consistency with previous MEG HMM studies^35,41^. Binary state time series of state activation/inactivation allowed to determine several state temporal parameters such as the MLT (mean duration of time intervals of active state), the FO (total fraction of time during which the state is active) and the MIL (mean duration of time intervals of inactive state). State power maps, which identify the topography of state-specific power envelope changes during state activation vs. during state inactivation, were computed as the partial correlation between states binary time series and group-concatenated power envelopes^35^.

### Statistical analyses

In order to discard possible confounds attributable to power, sex and version of the MEG system used in this study (either Vectorview or Triux), these parameters were regressed out of static rsFC prior to statistical analysis. For investigation of changes in power and in state temporal parameters (i.e., MLT, FO, MIL), only sex and MEG system version were regressed out. Statistical differences between groups (i.e., children, young adults and elders) were assessed using non-parametric Kruskal-Wallis tests with post-hoc Tukey’s range test on ranks to disclose age-related effects. For global connectivity and global power, significance was set at *p* < 0.05 Bonferroni corrected for the number of frequency bands, whereas for mean network connectivity and mean network power, significance was set at *p* < 0.05 Bonferroni corrected for the number of frequency bands times the number of RSNs. For static rsFC connectomes, significance was set at *p* < 0.05 Bonferroni corrected for the number of effective band-specific connections. The latter was directly assessed using the number of spatial degrees of freedom (ρ) estimated from the rank of the leadfield^16,54^, here ρ=55. Taking into account the symmetry of rsFC matrices and number of frequency bands investigated, the number of effective band-specific connections was *N*_eff_ = 3 × ρ × (ρ–1)/2 ≈ 513. To determine the proportion of significant within-RSN (respectively, cross-RSNs) connections, we divided the number of significant within-RSN (respectively, cross-RSNs) connections (summed over all RSNs) by the total number of possible within-RSN (respectively, cross-RSNs) connections. Finally, for temporal parameters associated to each transient HMM state, significance level was similarly set at *p* < 0.05 Bonferroni corrected for the number of independent states (i.e., *K*-1=7; the loss of one degree of freedom being due to the model constraint that one and only only state is active at any given time point). The state power maps were also statistically thresholded using a two-tailed parametric correlation tests at *p* < 0.05. The null hypothesis tested was that Fisher-transformed correlations follow a Gaussian distribution with mean zero and SD 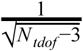. The number *N* of temporal degrees of freedom was estimated as one-quarter of the total number of time samples in group-concatenated envelope signals at 40 Hz sampling frequency, to take into account the 75% overlap in the envelope downsampling. The critical *p*-value was Bonferroni corrected with the number of independent HMM states (*K*-1=7) times the number of spatial degrees of freedom (ρ=55). Positive values greater than the significance level were considered as significant and disclosed regions with significant power increase/decrease upon state activation/inactivation. Respectively, negative values below the opposite of the significance level were considered as significant and identified regions with significant power decrease/increase upon state activation/inactivation.

### Data availability

The datasets analysed in this study are available from the corresponding author on reasonable request.

## Acknowledgments

This study was supported by the Action de Recherche Concertée Consolidation (ARCC, **“**Characterizing the spatio-temporal dynamics and the electrophysiological bases of resting state networks**”**, ULB, Brussels, Belgium), the Fonds Erasme (Research Convention **“**Les Voies du Savoir**”**,Brussels, Belgium) and the Fonds de la Recherche Scientifique (Research Convention: T.0109.13, FRS-FNRS, Brussels, Belgium). Nicolas Coquelet has been supported by the ARCC, by the Fonds Erasme (Research Convention **“**Les Voies du Savoir**”**, Brussels, Belgium) and is supported by the FRS-FNRS (Research Convention: Excellence of Science EOS “MEMODYN”). Alison Mary is Postdoctoral Researcher at the FRS-FNRS. Maxime Niesen and Marc Vander Ghinst have been supported by the Fonds Erasme. Mathieu Bourguignon is supported by the program Attract of Innoviris (Research Grant 2015-BB2B-10, Brussels, Belgium), the Marie Sklodowska-Curie Action of the European Commission (Research Grant: 743562) and by the Spanish Ministery of Economy and Competitiveness (Research Grant: PSI2016-77175-P). Xavier De Tiège is Postdoctorate Clinical Master Specialist at the FRS-FNRS.

The MEG project at the CUB Hôpital Erasme is financially supported by the Fonds Erasme.

## Author contributions

N.C., V.W. and X.D.T. designed study; N.C., V.W., A.M., M.N., D.P., M.R. and M.V.G. acquired data; N.C. and V.W. contributed to analysis tools; N.C., V.W. and X.D.T. analysed data; N.C., V.W., A.M., M.N., D.P., M.R., M.V.G., M.B., P.P., S.G., M.W. and X.D.T. wrote and reviewed the manuscript.

## Additional Information

The authors declare that they have no competing financial interests.

## References

1. Vanderhaeghen, P. & Cheng, H.-J. Guidance molecules in axon pruning and cell death. Cold Spring Harb. Perspect. Biol. 2, a001859 (2010).

2. Stiles, J. Brain development and the nature versus nurture debate. Prog. Brain Res. 189, 3–22 (2011).

3. Casey, B. J., Galvan, A. & Hare, T. A. Changes in cerebral functional organization during cognitive development. Current Opinion in Neurobiology. 15, 239–244 (2005).

4. Grayson, D. S. & Fair, D. A. Development of large-scale functional networks from birth to adulthood: A guide to the neuroimaging literature. Neuroimage. 160, 15–31 (2017).

5. Khundrakpam, B. S., Lewis, J. D., Zhao, L., Chouinard-Decorte, F. & Evans, A. C. Brain connectivity in normally developing children and adolescents. Neuroimage. 134, 192–203 (2016).

6. Ernst, M., Torrisi, S., Balderston, N., Grillon, C. & Hale, E. A. fMRI functional connectivity applied to adolescent neurodevelopment. Annu. Rev. Clin. Psychol. 11, 361–377 (2015).

7. Oldham, S. & Fornito, A. The Development of Brain Network Hubs. Dev. Cogn. Neurosci. 36, 100607 (2019).

8. Fjell, A. M., McEvoy, L., Holland, D., Dale, A. M. & Walhovd, K. B. What is normal in normal aging? Effects of aging, amyloid and Alzheimer’s disease on the cerebral cortex and the hippocampus. Progress in Neurobiology. 117, 20–40 (2014).

9. Zuo, X.-N. et al. Human Connectomics across the Life Span. Trends Cogn. Sci. 21, 32–45 (2017).

10. Bano, D., Agostini, M., Melino, G. & Nicotera, P. Ageing, neuronal connectivity and brain disorders: an unsolved ripple effect. Mol. Neurobiol. 43, 124–130 (2011).

11. Damoiseaux, J. S. Effects of aging on functional and structural brain connectivity. NeuroImage. 160, 32–40 (2017).

12. Toepper, M. Dissociating Normal Aging from Alzheimer’s Disease: A View from Cognitive Neuroscience. Journal of Alzheimer’s Disease. 57, 331–352 (2017).

13. Sala-Llonch, R., Bartrés-Faz, D. & Junqué, C. Reorganization of brain networks in aging: a review of functional connectivity studies. Front. Psychol. 6, 663 (2015).

14. Salthouse, T. A. Neuroanatomical substrates of age-related cognitive decline. Psychol. Bull. 137, 753–784 (2011).

15. Sun, F. W. et al. Youthful Brains in Older Adults: Preserved Neuroanatomy in the Default Mode and Salience Networks Contributes to Youthful Memory in Superaging. J. Neurosci. 36, 9659–9668 (2016).

16. Coquelet, N. et al. The electrophysiological connectome is maintained in healthy elders: a power envelope correlation MEG study. Scientific Reports. 7 (2017).

17. Trotta, N. et al. Functional integration changes in regional brain glucose metabolism from childhood to adulthood. Hum. Brain Mapp. 37, 3017–3030 (2016).

18. Cabeza, R. et al. Age-related changes in neural interactions during memory encoding and retrieval: A network analysis of PET data. Brain Cogn. 35, 369–372 (1997).

19. Nagel, I. E. et al. Performance level modulates adult age differences in brain activation during spatial working memory. Proc. Natl. Acad. Sci. U. S. A. 106, 22552–22557 (2009).

20. Luna, B., Padmanabhan, A. & O’Hearn, K. What has fMRI told us about the development of cognitive control through adolescence? Brain Cogn. 72, 101–113 (2010).

21. Fox, M. D. & Raichle, M. E. Spontaneous fluctuations in brain activity observed with functional magnetic resonance imaging. Nature Reviews Neuroscience. 8, 700–711 (2007).

22. Raichle, M. E. Two views of brain function. Trends in Cognitive Sciences. 14, 180–190 (2010).

23. Deco, G. & Corbetta, M. The Dynamical Balance of the Brain at Rest. The Neuroscientist. 17, 107–123 (2011).

24. Brookes, M. J. et al. Investigating the electrophysiological basis of resting state networks using magnetoencephalography. Proc. Natl. Acad. Sci. U. S. A. 108, 16783–16788 (2011).

25. de Pasquale, F. et al. Temporal dynamics of spontaneous MEG activity in brain networks. Proc. Natl. Acad. Sci. U. S. A. 107, 6040–6045 (2010).

26. Hipp, J. F., Hawellek, D. J., Corbetta, M., Siegel, M. & Engel, A. K. Large-scale cortical correlation structure of spontaneous oscillatory activity. Nat. Neurosci. 15, 884–890 (2012).

27. Wens, V. et al. About the electrophysiological basis of resting state networks. Clin. Neurophysiol. 125, 1711–1713 (2014).

28. Brookes, M. J. et al. Measuring temporal, spectral and spatial changes in electrophysiological brain network connectivity. Neuroimage 91, 282–299 (2014).

29. Liu, Q., Ganzetti, M., Wenderoth, N. & Mantini, D. Detecting Large-Scale Brain Networks Using EEG: Impact of Electrode Density, Head Modeling and Source Localization. Front. Neuroinform. 12, 4 (2018).

30. Liu, Q., Farahibozorg, S., Porcaro, C., Wenderoth, N. & Mantini, D. Detecting large-scale networks in the human brain using high-density electroencephalography. Hum. Brain Mapp. 38, 4631–4643 (2017).

31. Siems, M., Pape, A.-A., Hipp, J. F. & Siegel, M. Measuring the cortical correlation structure of spontaneous oscillatory activity with EEG and MEG. Neuroimage. 129, 345–355 (2016).

32. Coquelet, N. et al. Comparing MEG and high-density EEG for intrinsic functional connectivity mapping. Neuroimage. 210, 116556 (2020).

33. Hari, R. & Puce, A. Data Acquisition and Preprocessing. MEG-EEG Primer 89–97 (2017) doi: 10.1093/med/9780190497774.003.0007.

34. Vidaurre, D. et al. Spectrally resolved fast transient brain states in electrophysiological data. Neuroimage. 126, 81–95 (2016).

35. Baker, A. P. et al. Fast transient networks in spontaneous human brain activity. Elife. 3, e01867 (2014).

36. Wens, V. et al. Synchrony, metastability, dynamic integration, and competition in the spontaneous functional connectivity of the human brain. Neuroimage. 199, 313–324 (2019).

37. D’Esposito, M., Deouell, L. Y. & Gazzaley, A. Alterations in the BOLD fMRI signal with ageing and disease: a challenge for neuroimaging. Nat. Rev. Neurosci. 4, 863–872 (2003).

38. Barkhof, F., Haller, S. & Rombouts, S. A. R. B. Resting-state functional MR imaging: a new window to the brain. Radiology. 272, 29–49 (2014).

39. Briley, P. M. et al. Development of human electrophysiological brain networks. J. Neurophysiol. 120, 3122–3130 (2018).

40. Schäfer, C. B., Morgan, B. R., Ye, A. X., Taylor, M. J. & Doesburg, S. M. Oscillations, networks, and their development: MEG connectivity changes with age. Hum. Brain Mapp. 35, 5249–5261 (2014).

41. Brookes, M. J. et al. Altered temporal stability in dynamic neural networks underlies connectivity changes in neurodevelopment. Neuroimage. 174, 563–575 (2018).

42. O’Neill, G. C. et al. Dynamics of large-scale electrophysiological networks: A technical review. Neuroimage. 180, 559–576 (2018).

43. Vidaurre, D. et al. Spontaneous cortical activity transiently organises into frequency specific phase-coupling networks. Nat. Commun. 9, 2987 (2018).

44. Sitnikova, T. A., Hughes, J. W., Ahlfors, S. P., Woolrich, M. W. & Salat, D. H. Short timescale abnormalities in the states of spontaneous synchrony in the functional neural networks in Alzheimer’s disease. Neuroimage Clin. 20, 128–152 (2018).

45. Hawkins, E. et al. Functional network dynamics in a neurodevelopmental disorder of known genetic origin. Hum. Brain Mapp. 41, 530–544 (2020).

46. Quinn, A. J. et al. Task-Evoked Dynamic Network Analysis Through Hidden Markov Modeling. Frontiers in Neuroscience. 12 (2018).

47. Sjøgård, M. et al. Do the posterior midline cortices belong to the electrophysiological default-mode network? Neuroimage. 200, 221–230 (2019).

48. Yang, Z. et al. Connectivity trajectory across lifespan differentiates the precuneus from the default network. Neuroimage. 89, 45–56 (2014).

49. Bagarinao, E. et al. Reorganization of brain networks and its association with general cognitive performance over the adult lifespan. Sci. Rep. 9, 11352 (2019).

50. de Pasquale, F. et al. A cortical core for dynamic integration of functional networks in the resting human brain. Neuron. 74, 753–764 (2012).

51. Brookes, M. J., Woolrich, M. W. & Barnes, G. R. Measuring functional connectivity in MEG: a multivariate approach insensitive to linear source leakage. Neuroimage. 63, 910–920 (2012).

52. Colclough, G. L., Brookes, M. J., Smith, S. M. & Woolrich, M. W. A symmetric multivariate leakage correction for MEG connectomes. Neuroimage. 117, 439–448 (2015).

53. Palva, J. M. et al. Ghost interactions in MEG/EEG source space: A note of caution on inter-areal coupling measures. Neuroimage. 173, 632–643 (2018).

54. Wens, V. et al. A geometric correction scheme for spatial leakage effects in MEG/EEG seed-based functional connectivity mapping. Hum. Brain Mapp. 36, 4604–4621 (2015).

55. Rezek, I. & Roberts, S. Ensemble Hidden Markov Models with Extended Observation Densities for Biosignal Analysis. in Probabilistic Modeling in Bioinformatics and Medical Informatics (eds. Husmeier, D., Dybowski, R. & Roberts, S.) vol. 16 419–450 (Springer-Verlag, 2005).

56. Hagmann, P. et al. White matter maturation reshapes structural connectivity in the late developing human brain. Proc. Natl. Acad. Sci. U. S. A. 107, 19067–19072 (2010).

57. Seghier, M. L. The angular gyrus: multiple functions and multiple subdivisions. Neuroscientist. 19, 43–61 (2013).

58. Igelström, K. M. & Graziano, M. S. A. The inferior parietal lobule and temporoparietal junction: A network perspective. Neuropsychologia. 105, 70–83 (2017).

59. Li, R. et al. Developmental Maturation of the Precuneus as a Functional Core of the Default Mode Network. J. Cogn. Neurosci. 31, 1506–1519 (2019).

60. Andrews-Hanna, J. R., Reidler, J. S., Sepulcre, J., Poulin, R. & Buckner, R. L. Functional-Anatomic Fractionation of the Brain’s Default Network. Neuron. 65, 550–562 (2010).

61. Uddin, L. Q., Kelly, A. M., Biswal, B. B., Castellanos, F. X. & Milham, M. P. Functional connectivity of default mode network components: correlation, anticorrelation, and causality. Hum. Brain Mapp. 30, 625–637 (2009).

62. Grady, C. L. et al. Age-related changes in cortical blood flow activation during visual processing of faces and location. J. Neurosci. 14, 1450–1462 (1994).

63. Park, D. C. et al. Aging reduces neural specialization in ventral visual cortex. Proc. Natl. Acad. Sci. U. S. A. 101, 13091–13095 (2004).

64. Zonneveld, H. I. et al. Patterns of functional connectivity in an aging population: The Rotterdam Study. NeuroImage. 189, 432–444 (2019).

65. Andrews-Hanna, J. R. et al. Disruption of large-scale brain systems in advanced aging. Neuron. 56, 924–935 (2007).

66. Tomasi, D. & Volkow, N. D. Aging and functional brain networks. Molecular Psychiatry. 17, 549–558 (2012).

67. Geerligs, L., Renken, R. J., Saliasi, E., Maurits, N. M. & Lorist, M. M. A Brain-Wide Study of Age-Related Changes in Functional Connectivity. Cereb. Cortex. 25, 1987–1999 (2015).

68. Kucyi, A., Tambini, A., Sadaghiani, S., Keilholz, S. & Cohen, J. R. Spontaneous cognitive processes and the behavioral validation of time-varying brain connectivity. Netw Neurosci 2, 397–417 (2018).

69. Stawarczyk, D., Majerus, S., Catale, C. & D’Argembeau, A. Relationships between mind-wandering and attentional control abilities in young adults and adolescents. Acta Psychol. 148, 25–36 (2014).

70. Keulers, E. H. H. & Jonkman, L. M. Mind wandering in children: Examining task-unrelated thoughts in computerized tasks and a classroom lesson, and the association with different executive functions. Journal of Experimental Child Psychology. 179, 276–290 (2019).

71. Zhang, Y., Song, X., Ye, Q. & Wang, Q. Children with positive attitudes towards mind-wandering provide invalid subjective reports of mind-wandering during an experimental task. Conscious. Cogn. 35, 136–142 (2015).

72. Ye, Q., Song, X., Zhang, Y. & Wang, Q. Children’s mental time travel during mind wandering. Frontiers in Psychology. 5 (2014).

73. Maillet, D. et al. Aging and the wandering brain: Age-related differences in the neural correlates of stimulus-independent thoughts. PLoS One. 14, e0223981 (2019).

74. McCormack, T., Burns, P., O’Connor, P., Jaroslawska, A. & Caruso, E. M. Do children and adolescents have a future-oriented bias? A developmental study of spontaneous and cued past and future thinking. Psychological Research. 83, 774–787 (2019).

75. Maillet, D. & Schacter, D. L. From mind wandering to involuntary retrieval: Age-related differences in spontaneous cognitive processes. Neuropsychologia. 80, 142–156 (2016).

76. Seli, P., Maillet, D., Smilek, D., Oakman, J. M. & Schacter, D. L. Cognitive aging and the distinction between intentional and unintentional mind wandering. Psychol. Aging. 32, 315–324 (2017).

77. Maillet, D. et al. Age-related differences in mind-wandering in daily life. Psychol. Aging. 33, 643–653 (2018).

78. Hill, R. M. et al. A tool for functional brain imaging with lifespan compliance. Nat. Commun. 10, 4785 (2019).

79. Rolls, E. T., Huang, C.-C., Lin, C.-P., Feng, J. & Joliot, M. Automated anatomical labelling atlas 3. Neuroimage. 206, 116189 (2020).

80. Yesavage, J. A. & Sheikh, J. I. 9/Geriatric Depression Scale (GDS): Recent Evidence and Development of a Shorter Version. Clin. Gerontol. 5, 165–173 (1986).

81. Morris, J. C. The Clinical Dementia Rating (CDR): current version and scoring rules. Neurology. 43, 2412–2414 (1993).

82. Folstein, M. F., Folstein, S. E. & McHugh, P. R. ‘Mini-mental state’. J. Psychiatr. Res. 12, 189–198 (1975).

83. Folstein, M. F., Robins, L. N. & Helzer, J. E. The Mini-Mental State Examination. Arch. Gen. Psychiatry. 40, 812 (1983).

84. Fastenau, P. S., Denburg, N. L. & Hufford, B. J. Adult Norms for the Rey-Osterrieth Complex Figure Test and for Supplemental Recognition and Matching Trials from the Extended Complex Figure Test. The Clinical Neuropsychologist. 13, 30–47 (1999).

85. Cardebat, D., Doyon, B., Puel, M., Goulet, P. & Joanette, Y. [Formal and semantic lexical evocation in normal subjects. Performance and dynamics of production as a function of sex, age and educational level]. Acta Neurol. Belg. 90, 207–217 (1990).

86. Baddeley, A. D., Emslie, H. & Nimmo-Smith, I. Doors and People: A Test of Visual and Verbal Recall and Recognition. Manual. (1994).

87. Wechsler, D. Wechsler memory scale-revised manual. (1987).

88. Van der Linden, M. et al. L’épreuve de rappel libre / rappel indicé à 16 items (RL/RI-16). (Solal, 2004).

89. Bachy Langedock, N. Batterie d’examen des troubles de la dénomination (ExaDé). (1988).

90. Tombaugh, T. Trail Making Test A and B: Normative data stratified by age and education. Archives of Clinical Neuropsychology. 19, 203–214 (2004).

91. MacLeod, C. M. Half a century of research on the Stroop effect: an integrative review. Psychol. Bull. 109, 163–203 (1991).

92. Tiège, X. D. et al. Recording epileptic activity with MEG in a light-weight magnetic shield. Epilepsy Research. 82, 227–231 (2008).

93. Marty, B. et al. Evidence for genetically determined degeneration of proprioceptive tracts in Friedreich ataxia. Neurology. 93, e116–e124 (2019).

94. Naeije, G. et al. Altered neocortical tactile but preserved auditory early change detection responses in Friedreich ataxia. Clin. Neurophysiol. 130, 1299–1310 (2019).

95. Naeije, G. et al. Age of onset determines intrinsic functional brain architecture in Friedreich ataxia. Ann Clin Transl Neurol. 7, 94–104 (2020).

96. Taulu, S., Simola, J. & Kajola, M. Applications of the signal space separation method. IEEE Trans. Signal Process. 53, 3359–3372 (2005).

97. Vigário, R., Särelä, J., Jousmäki, V., Hämäläinen, M. & Oja, E. Independent component approach to the analysis of EEG and MEG recordings. IEEE Trans. Biomed. Eng. 47, 589–593 (2000).

98. Dale, A. M. & Sereno, M. I. Improved Localizadon of Cortical Activity by Combining EEG and MEG with MRI Cortical Surface Reconstruction: A Linear Approach. Journal of Cognitive Neuroscience. 5, 162–176 (1993).

99. Woolrich, M. W. et al. Dynamic state allocation for MEG source reconstruction. Neuroimage. 77, 77–92 (2013).

